# Latitudinal gradients in predation persist in urban environments

**DOI:** 10.1101/2023.11.14.566324

**Authors:** Anna L Hargreaves, John Ensing, Olivia Rahn, Fernanda M. P. Oliveira, Jérôme Burkiewicz, Joëlle Lafond, Sybille Haeussler, M. Brooke Byerley-Best, Kira Lazda, Heather L. Slinn, Ella Martin, Matthew L. Carlson, Todd L. Sformo, Emma Dawson-Glass, Mariana C. Chiuffo, Yalma L. Vargas-Rodriguez, Carlos I. García-Jiménez, Inácio J. M. T. Gomes, Sandra Klemet-N’Guessan, Lucas Paolucci, Simon Joly, Klaus Mehltreter, Jenny Muñoz, Carmela Buono, Jedediah F. Brodie, Antonio Rodriguez-Campbell, Thor Veen, Ben Freeman, Julie Lee-Yaw, Juan Camilo Muñoz, Alexandra Paquette, Jennifer Butler, Esteban Suaréz

**Affiliations:** Department of Biology, McGill University; Montreal, QC, Canada; Department of Biology, Okanagan College; Vernon, BC, Canada; Departamento de Ciências Biológica, Universidade de Pernambuco; Campus Garanhuns, Pernambuco, Brasil; Département de Sciences Biologiques, Université de Montréal; Montréal, QC, Canada; University of Northern British Columbia; Smithers, BC, Canada; Botanical Research Institute of Texas, Fort Worth Botanic Garden; Fort Worth, TX, USA; Integrative Biology, University of Guelph; Guelph, ON, Canada; Alaska Center for Conservation Science, University of Alaska; Anchorage, AK, USA; Institute of Arctic Biology, University of Alaska; Fairbanks, AK, USA; INIBIOMA, Universidad Nacional del Comahue, CONICET; San Carlos de Bariloche, Río Negro, Argentina; Centro Universitario de los Valles, Universidad de Guadalajara; Ameca, Jalisco, México; Universidad de Guadalajara; Zapopan, Jalisco, México; Departamento de Biologia Geral, Universidade Federal de Viçosa; Viçosa, Minas Gerais, Brazil; Instituto de Ciências Biológicas, Universidade Federal de Minas Gerais; Belo Horizonte, Minas Gerais, Brazil; Department of Biology, Trent University; Peterborough, ON, Canada; Montreal Botanical Garden, Montreal, QC, Canada; Red de Ecología Funcional, Instituto de Ecología, Xalapa, Veracruz, México; Biodiversity Research Centre, University of British Columbia, Vancouver, BC, Canada; Department of Biological Sciences, SUNY Binghamton University, Binghamton, NY, USA; Biological Science & Wildlife Biology Program, University of Montana, Missoula, MT, USA; Institute for Biodiversity & Environmental Conservation, Universiti Malaysia Sarawak, Kota Samarahan, Sarawak, Malaysia; Quest University Canada, Squamish, BC, Canada; Department of Biological Sciences, University of Lethbridge, Lethbridge, AB, Canada; Fundación Humedales, Bogotá, Colombia; Institute for Applied Ecology, Corvallis, OR, USA; Instituto Biósfera & Colegio de Ciencias Biológicas y Ambientales, Universidad San Francisco de Quito, Ecuador

## Abstract

Urbanization can profoundly disrupt local ecology. But while urban areas now stretch across latitudes, little is known about urbanization’s effects on macroecological patterns. We used standardized experiments to test whether urbanization disrupts latitudinal gradients in seed predation, a macroecological pattern that shapes community assembly and diversity. Using >56,000 seeds, we compared predation in urbanized and natural areas across 14,000 km of latitude, spanning the Americas. Predation increased 5-fold from high latitudes to the tropics, and latitudinal gradients in predation persisted in urban areas despite significant habitat modification. Urbanization reduced predation by vertebrates, but not invertebrates, and seemed to increase ant predation specifically. Our results show that macroecological patterns in predation intensity can persist in urbanized environments, even as urbanization alters the relative importance of predators.

**One-Sentence Summary:** Across 56,000 seeds and 112° of latitude, latitudinal gradients in seed predation are equally strong in natural vs. urban areas

## Introduction

Biologists have long speculated that interactions among species become stronger toward the tropics (*1*). This latitudinal gradient in increasing interaction intensity, paralleling gradients in temperature, primary productivity and biodiversity, is thought to have played a major role in shaping global patterns in ecology and evolution (*2–4*). For example, increasing risk of being eaten from high to low latitudes is invoked to explain long-distance migration (*5*), species’ equatorward range edges (*6*), the astounding diversity of tropical forests (*7, 8*), and faster speciation in the tropics (*9*). Large-scale experiments have now demonstrated dramatic latitudinal gradients in the intensity of predation, both within and across terrestrial and coastal biomes (*5, 10–12*). However, such experiments have so far been restricted to large natural areas; it remains unclear how robust these macroecological patterns are to accelerating anthropogenic change.

One of the most intense and rapidly expanding forms of land use change is urbanization (*13, 14*). Its ecological effects can be seen even at its earliest stages, as natural vegetation in rural areas is cleared to make way for homes (*15*). As urbanization intensifies, most species decline in abundance or disappear altogether (*16, 17*) though a few experience population booms (*18, 19*). This biotic reshuffling can alter the nature and intensity of local trophic interactions (*20–22*), impacting ecosystem services, ecological functioning, and evolution in cities and their suburbs (*23–25*). But urbanization is more than a patchwork of local effects; it has created a novel biome in which urbanized areas often resemble other, distant urbanized areas more than adjacent natural areas (*26, 27*). While local effects of urbanization are well studied, we know little about its effects on macroecological patterns (*28, 29*).

Here, we test whether the homogenizing effect of urbanization flattens latitudinal gradients in predation intensity. We consider post-dispersal seed predation, a global interaction that shapes the structure and distribution of plant communities (*30, 31*). Seed predation is particularly relevant to theory about latitudinal gradients in species interactions as increased seed predation toward the tropics is thought to both ecologically maintain and evolutionarily accelerate the spectacular diversity of tropical plants and their enemies (*7, 8, 32*). As the urban biome is set to double in size by 2100 (*33*), its effects on the macroecological patterns that shape modern biodiversity, such as latitudinal gradients in predation, are increasingly important to the future distribution and evolution of life.

One might expect urbanization to flatten latitudinal gradients by reducing latitudinal differences among habitats. Grassy lawns in Brazil and Canada resemble each other far more than the tropical and boreal forest they replaced (‘biotic homogenization’; Fig. S1). Indeed, both abiotic conditions (*15*) and species assemblages (*34, 35*) are more similar among urban areas than rural ones. Early evidence suggests that urbanization can flatten latitudinal gradients in species diversity (*29, 36*), which are proposed to both drive and reflect latitudinal gradients in species interactions (*2*). On the other hand, seed predation rates are only loosely related to predator diversity (*12, 37*), and more directly reflect predator abundance or activity. If urban predator assemblages maintain overall abundance or foraging intensity (*23, 38, 39*), urbanization may not affect predation rates even when it reduces biodiversity. Or, if urbanization reduces predator abundance similarly across latitudes, urbanized sites may have lower predation but still maintain latitudinal gradients in predation intensity.

Using consistent experimental methods and > 56,000 seeds, we compared latitudinal gradients in predation between urbanized sites (urban and suburban backyards) versus nearby natural areas across 112° of latitude (Fig. 1). This work is an extension of the Biotic Interaction Gradients (BIG) experiments, which found steep latitudinal clines in seed predation when measured in large wilderness areas (*12*); here we test whether these gradients are maintained in smaller natural areas and urbanized areas. To measure underlying predation rates, unclouded by local adaptations between seeds and predators, we used agricultural seeds with minimal chemical or physical defenses. We distinguished among three types of predation, as urbanization effects vary dramatically among taxa (*40*): total predation, measured using sunflower seeds without shells, which are eaten by diverse invertebrates and vertebrates; vertebrate predation using oats, which are eaten almost exclusively by small mammals and birds; and invertebrate predation by caging some sunflower seeds to exclude vertebrates (Figs S1&S2). We set seeds out in small piles (‘depots’) on the ground, in areas with vegetated ground cover, be it lawns, garden beds, or untended areas with natural litter. After 24 h we quantified predation and noted any indicators of predator identity (e.g. invertebrates still eating seeds, rodent feces).

**Fig. 1.**
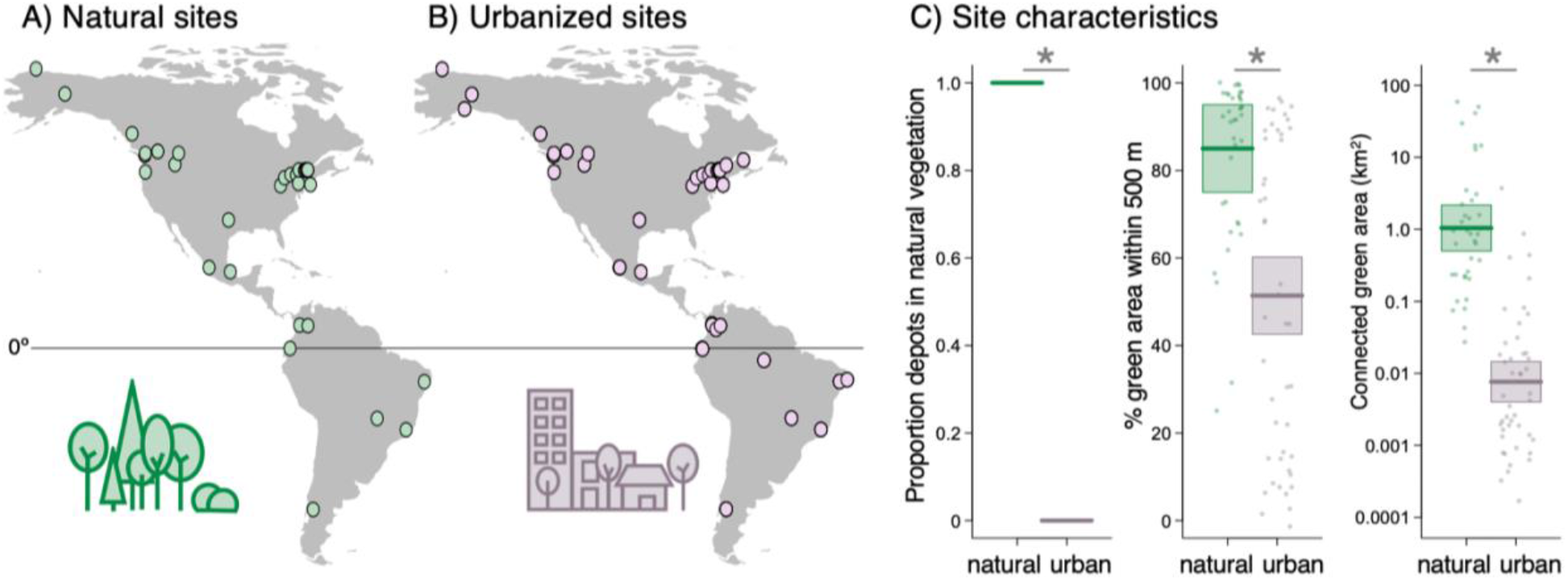
Site locations and characteristics. Locations of experimental sites in natural areas (A; *n* = 36) and in urbanized (backyard) areas (B; *n* = 45). C) Urbanized sites had less ground surface with untended, natural vegetation and litter (leftmost plot), less green area within a 500 m radius, and smaller connected green areas than natural sites (rightmost plot; green areas were defined as those not covered with buildings, paving or stonework). Horizontal lines, boxes and points show the mean, 95% CI, and partial residuals extracted from models with fixed effects Urbanization + absolute Latitude. None of the three response variables varied with latitude (statistical results in Table S1), so all are plotted for the median latitude (45°). CI and residuals in leftmost plot are too narrowly distributed around the mean to be visible.

## Results & Discussion

We found strong latitudinal gradients in seed predation, but no evidence that these gradients are flattened by urbanization. Across 337 runs of the experiment at 81 study sites, seed predation increased more than 5-fold from the poles to equator (Fig. 2). Latitudinal gradients in predation were as steep in urbanized areas as in natural areas (latitude × urbanization: χ^2^_df=1_ = 0.11, *P* = 0.74; Fig. 2), and this was true for total seed predation, predation by invertebrates only, and predation by vertebrates (latitude × urbanization × predation type: χ^2^_df=2_ = 1.21, *P* = 0.55). Results were consistent if we restricted data to the 303 runs where urbanized sites were tested with a concurrent paired natural site, and whether we modelled predation types together or separately (Table S2). Urbanized sites in our study had significantly less area with untended vegetation, lower local greenness, and smaller connected green areas than natural sites (Fig. 1C; Table S1). Thus our results provide robust evidence that despite significant habitat modification, global macroecological patterns can be maintained in urbanized areas.

**Fig. 2.**
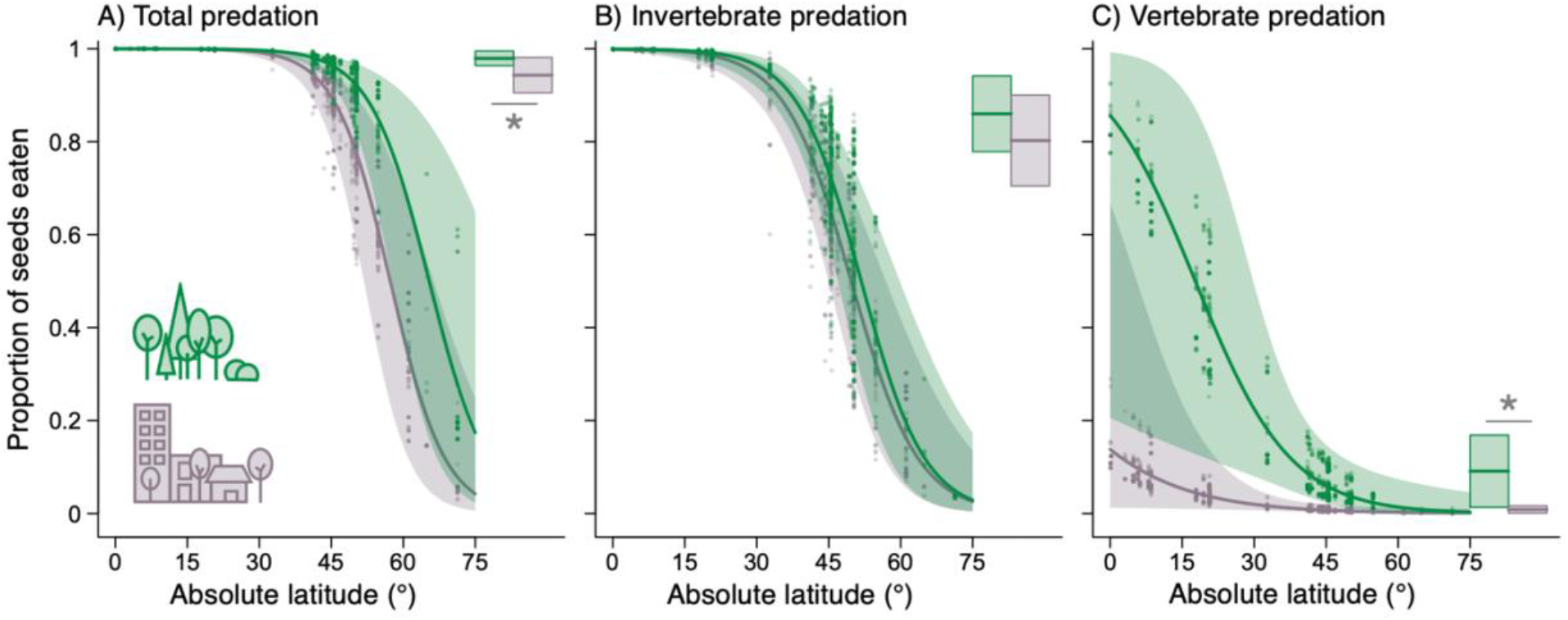
Latitudinal gradients in predation in natural and urbanized areas. Seed predation declined significantly toward higher latitudes for all predation types. Latitudinal patterns did not vary with urbanization for any predator group, i.e. green (higher) and purple (lower) lines do not differ significantly in slope for any panel (the apparently differing slopes in C are visually exaggerated due to back transformation from the logit scale). Bars show predation intensity (±95% CI) for the median latitude in our study (same y-axis as trend lines); total and vertebrate predation were significantly lower in urbanized (right, purple bars) versus natural areas (left, green bars; *s). Trends, means, 95% CI, and partial residuals were extracted from one binomial GLMM per predation type to show the independent effects of urbanization on each predator group (results from overall model shown in Fig S1). All results are shown for the median elevation (450 masl). Sample sizes: 156 experimental runs at 36 natural sites and 181 runs at 45 urbanized sites, with 2912, 2394, and 2718 seed depots measuring total, invertebrate, and vertebrate predation, respectively.

While latitudinal gradients in predation were maintained in urbanized areas, predation intensity was not. Urbanization reduced overall predation on seeds (Fig. 2A), though this effect was not uniform across predators (urbanization × predation type: χ^2^_df=2_ = 18.2, *P* = 0.0001). Urbanization had little effect on predation by invertebrates (Fig. 2B), but significantly reduced predation by vertebrates (Fig. 2C). Changes in the intensity and agents of predation can alter the selective regime experienced by prey and thus the evolutionary trajectory of urban populations (*41*), as different predator guilds impose differing selection on seed characteristics (*42*). Large-scale effects of urbanization on predator-driven selection have been inferred from differences in the relative abundance of consumers (*43*) and prevalence of anti-predator/herbivore defenses (*15, 44*) in urban versus natural areas. Our results provide large-scale evidence for a mechanistic link: urbanization changes the relative intensity of predation by different predator groups across latitudes.

Altered predation in urbanized areas was not surprising, but the nature of effects was unexpected. One might predict urbanization to reduce invertebrate predation, contrary to our results, since urbanization can strongly reduce invertebrate diversity and abundance (*17, 19, 45*). In particular, ants were our most commonly detected seed predators (Table S3) and urbanization reduces ant diversity, especially in the tropics (*29, 46*). However, across latitudes we observed ant predation *more* often in urbanized than natural areas (Fig. 3). So either granivorous ants are not suffering the same urbanization-driven declines in diversity as ants in general (*29*), or their abundance and activity are decoupled from diversity. Indeed, our results suggest that any lost diversity of seed-eating invertebrates in urbanized areas is being made up for by increased abundance or activity of remaining species (*47*). This result begs the more general question of how well diversity predicts species interaction intensity. While latitudinal gradients in diversity inspired the original hypotheses for latitudinal gradients in interaction intensity (*3*), testing this link has been challenging as diversity covaries with other potential drivers of interaction intensity (e.g. temperature, productivity (*11, 12, 37*)). By decoupling local diversity from latitude and climate (*36*), urbanization offers an tantalizing opportunity to disentangle the potential drivers of predation intensity.

**Fig. 3.**
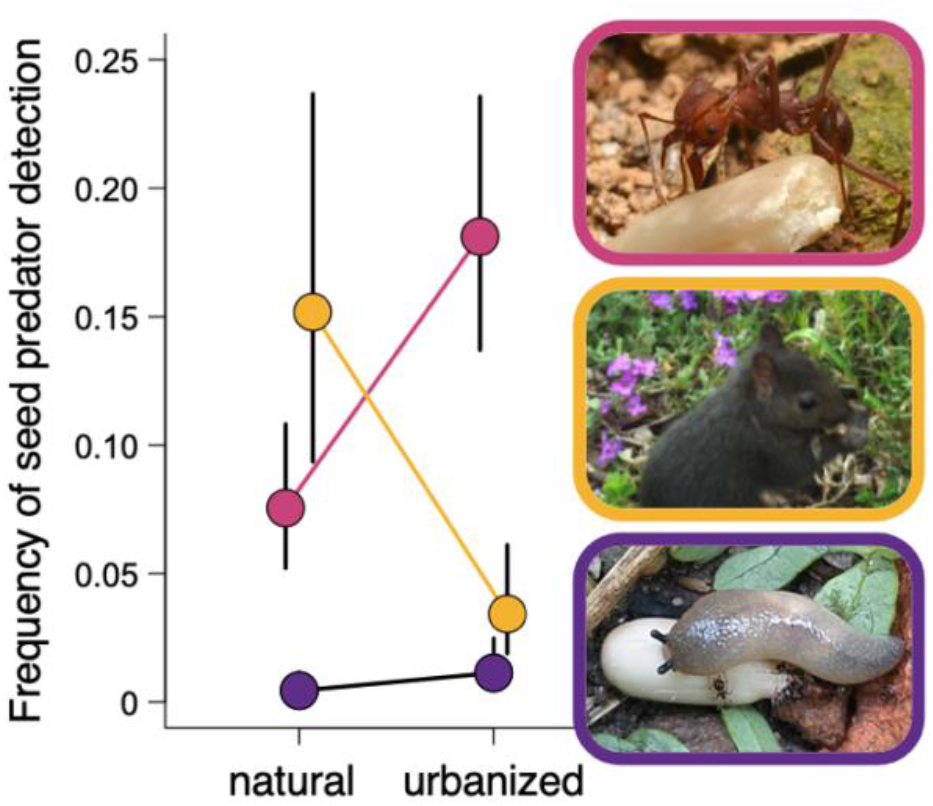
Effect of urbanization on observed seed predators. Of the 5829 seed depots in which seeds were eaten, we were able to identify the seed predators for 1469. For the three most commonly identified seed predators we tested whether their predation was detected at a higher frequency (i.e. at more seed depots) in urbanized or natural sites. Ant predation was more common in urbanized sites (pink; χ^2^_df=2_ = 13.2, *P* = 0.0003), whereas small mammal predation was more frequently detected in natural sites (light orange: χ^2^_df=2_ = 15.3, *P* < 0.0001; oat depots). Predation by molluscs (snails and slugs) was equally common in urbanized and natural sites (dark purple; χ^2^_df=2_ = 2.8, *P* = 0.094). Points are mean proportion of depots at which the predator was detected (actual predation rates of all three predators are probably higher), plotted for the median latitude and elevation, ±95% CI. Full statistical details in Table S3. Photos from top: leaf cutter ant consuming sunflower seed (Brazil, 21°S; L Paolucci); eastern grey squirrel eating oats (Canada, 44°N; A Hargreaves); slug eating sunflower seed (Mexico, 21°N, C García-Jiménez).

Conversely, given that small mammals and generalist birds can thrive in urbanized areas (*48–50*), reduced predation by vertebrates was surprising. In particular, we detected signs of small mammal predation significantly less often in urbanized vs. natural areas (Fig. 3). It is possible that vertebrate seed predators are still abundant in urbanized areas, but seeking higher-reward foods (*51*) and ignoring experimental seeds. However, it is unclear why richer urban food supply would not also distract invertebrates from experimental seeds, and why it would not eventually lead to increased seed predator abundance, resulting in spill-over predation on experimental seeds. Alternatively, our findings could reflect negative impacts of urbanization on the abundance or foraging activity of granivorous birds and mammals (*52*) across latitudes.

Urbanized sites differed from natural sites in both seed predation (Fig. 2) and habitat modification (Fig. 1C), but also varied in how urban they were. As the ecological impact of urbanization often varies with the amount and type of green space remaining (*16, 19, 48*), we tested whether more direct measures of local greenness better predicted seed predation intensity. The percentage of greenspace within a 500 m radius of each site and the total area of greenspace connected to the site explained invertebrate predation slightly better than simply classifying a site as urbanized or natural (slight improvements in marginal *R*^2^ for GLMMs and AIC) but explained vertebrate and total predation slightly worse (Table S4). While we may have under-sampled the least green urban areas (e.g. our focus on residential areas excluded fully paved industrial zones), the lack of relationship between local greenness and seed predation suggests that increased sampling of highly-urbanized areas would not alter our main results.

While we did not find an effect of local greenness, within urbanized sites predation did vary with local ground cover. Invertebrate predation was lowest for seeds placed in untended areas with natural vegetation and litter, whereas predation by vertebrates was lowest on lawns (ground vegetation × predation type: χ^2^_df=4_ = 16.2, *P* = 0.0028), perhaps due to lack of overhead cover from predators (*53*). Total predation was therefore higher in garden beds than in areas of lawn or natural vegetation litter, which somewhat contrasts the larger scale pattern of higher total predation in natural sites. Thus the factors that determine predation intensity at small scales (within urbanized sites) must differ from those acting at larger scales (between urbanized and natural sites) (*54*).

Finally, our study highlights the robustness of geographic gradients in seed predation. In contrast to our prior work testing latitudinal and elevational gradients in predation (*12*), our current sites vary much more in longitude, hemisphere, and the size and connectivity of natural areas, yet we recover equally strong and highly consistent patterns. Across our 36 natural sites, a seed’s chance of being eaten increased by 10% for every 10° decline in latitude; remarkably similar to the latitudinal cline found across 79 different sites in different years (*12*). Ant predation increased strongly toward lower latitudes, whereas mammal predation did not (Table S3), consistent with previous conclusions that invertebrates drive latitudinal gradients in predation intensity (*12*). While our current study was not designed to test elevational effects, seed predation increased strongly from high to low elevations in all analyses (Table S2; Fig. S4). This is the first time such a large distributed experiment measuring interaction strength has been repeated. Our work demonstrates remarkable consistency in large-scale gradients in seed predation across time and space, and in the face of extensive habitat modification from urbanization. That some macroecological patterns are maintained in urbanized areas, even as other patterns are not (*36*), provides a fertile ground for disentangling the mechanistic causes of the global patterns that shape our current and future biodiversity.

## Acknowledgements

We thank Paula Morales, Margot MacLaren, Vida Javidi, Pascale Caissy, Christa Mulder, Phillida Hargreaves, and Fabricio Baccaro for contributing experimental runs; J. Lane, M. Moffat, J. Best, P. Gomes, B Ricci, and A. Iglesias for help with fieldwork; and Érablière Charbonneau, Érablière Meunier, C Ostiguy, E&M Lafond, Gault Nature Reserve for access to sampling sites. **Funding**: the primary funding for this BIG experiment was from an NSERC discovery grant, McGill Liber Ero Conservation Fund grant, and NFRF-R grant to A.L.H. We are also grateful for support from an NSERC graduate scholarship (E.M.); MITACs award (A.R.C.); NSERC and McGill undergraduate summer research awards (K.L., A.P., V.Javidi, M.Maclaren); and FONCyT (PICT 2019 00969) and CONICET (PIBAA 2022-2023) to M.Chiuffo. **Author contributions**: Authors are listed in order of contribution. A.L.H. designed and coordinated the study, analyzed the data, and wrote the manuscript. All authors contributed fieldwork, except O.R.; O.R. and K.L. developed greenness methods and collected greenness data. E.D.G. made cages and shipped standardized seeds to collaborators in USA. O.R., J.L., S.H., M.B.B, E.M, M.L.C, T.L.S., M.C.C., I.J.M.TG., S.K.N, L.P., S.J, K.M., J.F.B., B.F., J.L., E.S. helped edit the manuscript. **Competing interests**: Authors declare no competing interests. **Data and materials availability**: R code and data used in analyses will be made publicly available on Borealis, the Canadian Dataverse Repository.

## Supplementary Materials

### Materials and Methods

#### Experimental Design

We began our experiment during the COVID pandemic lockdowns of 2020, designing it as one we could conduct while confined to our backyards and local natural areas. Data were collected from 2020 to 2023. We included sites anywhere on the mainland of the Americas, excluding islands as they experience unique biogeographic processes that might obscure latitudinal patterns, and made sure to have good representation in the tropics and high latitudes. In total, we ran the experiment in eight countries spanning 112° of latitude.

Each collaborator measured seed predation in an urbanized site and, ideally, a paired natural area (Fig. S1; some collaborators ran the experiment in additional urbanized or natural sites as well). Urbanized sites were generally backyards, defined loosely as the unpaved (no asphalt, gravel, stonework etc.) area around a home, be it an apartment building, row house, detached house, or university residence, but in two cases were urban parks or vacant lots. As we only placed seed depots on unpaved ground, urbanized sites had to have some such areas (e.g. green balconies or cobblestone courtyards would be ineligible). Urbanized sites were thus unified in that they: 1) were all beside built structures continuously occupied by people (homes but also stores and other buildings in some cases); 2) were close to driveways and roads which separated them from other greenspace; and 3) had their natural vegetation cleared to make way for homes, replaced with modified vegetation (e.g. mowed lawns, garden beds). Some of these sites were highly urban (e.g. downtown Montreal or Guadalajara, <5% of surrounding area was vegetated), while others were in much greener areas. For this reason we use the term ‘urbanized’ rather than ‘urban’ throughout. Paired natural sites had the natural vegetation that would have presumably occupied the urbanized site prior to vegetation clearing, at roughly the same elevation.

We ran the experiment 1 to 11 times per site, pairing runs between urbanized and natural sites within a few days of each other when possible (each ‘run’ is one 24 h assay of predation at 1 site). Replicate runs within a site were separated by at least 2 weeks. Experiments were conducted during the snow-free growing season at each site, during weather typical for that site in that season.

**Fig S1.**
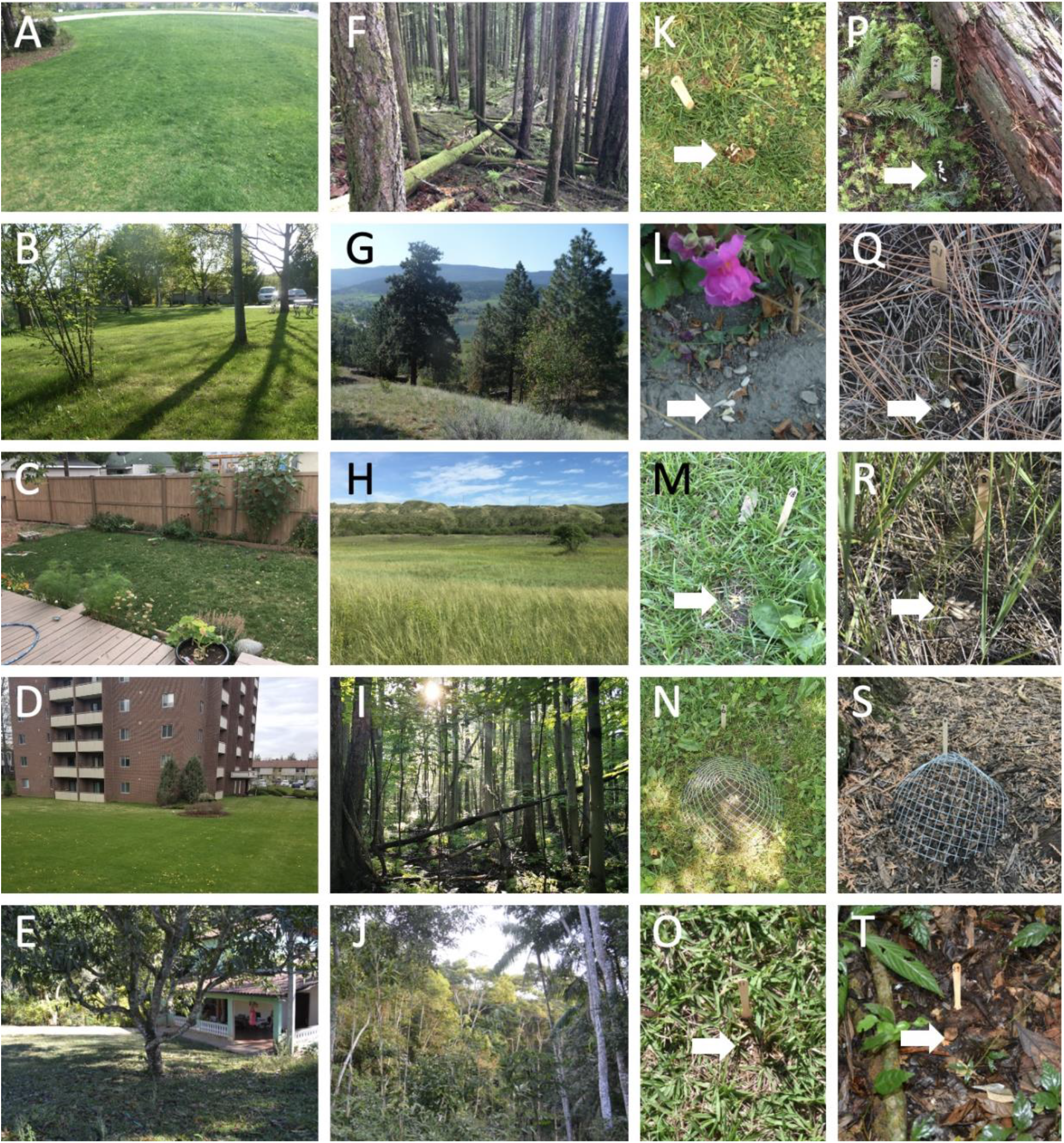
Experimental sites. Urbanized sites (A-E) often appear more similar to each other than natural sites (F-J), and often share more similar ground vegetation (K-O) than natural sites (P-T). By row: row A) coastal BC (50°N; wet conifer forest biome); row B) interior BC (50°N; dry conifer-grassland biome); row C) southern Alberta (50°N; prairie biome); row D) Great lakes-St Lawrence lowlands, Canada (44°N; mixed hardwood forest biome); row E) southeastern Brazil (21°S; tropical rainforest biome). Arrows indicate depots; vertebrate exclusion cages shown in N&S. Vegetation types: KMNO=lawn, L=garden, P-T=natural litter. Photo credits: AFKP=T Veen; BGLQ=J Ensing; CHMR=J Lee Yaw; D=H Slinn; IN=J Lafond; EJOR=L Paolucci; S S Klemet-N’Guessan.

#### Quantifying seed predation

This is an extension of the Biotic Interaction Gradients (BIG) experiment. As in previous BIG experiments (*12, 37*), at each site we set out small piles (‘depots’) of seeds directly on the ground (Fig. S1K-T). We deployed up to 30 depots per site per run as space allowed, separated by at least 6 m (mean = 22.6 depots per run at urbanized sites, 25.3 per run at natural sites). We returned after 24 h to quantify post-dispersal seed predation. To measure underlying predation rates, unclouded by local adaptations between seeds and predators, we used agricultural seeds bred for human-consumption; such seeds are not local to any site and should be widely palatable. We only used intact seeds so that damage was unambiguously be attributable to granivores. We used sunflower seeds with shells removed (8 seeds per depot; e.g. Fig S1. L) and oat seeds with their thin hulls intact (5 seeds per depot, e.g. Fig. S1Q).

We are familiar with the predators of sunflower and oat seeds from direct observations during depot-checks (at 24 h and sometimes earlier checks assessing how quickly seeds were eaten), opportunistic camera-trap observations, and ad-hoc feeding trials by A.L.H. during this and previous BIG experiments. We know that oil-based sunflower seeds are eaten by diverse predators, including small mammals (e.g. mice, voles or shrews (detected via feces), rats (camera traps), squirrels (camera traps)); birds (e.g. jays, robins, sparrows, finches (camera traps)); and a diverse array of invertebrates including ants, slugs, snails, beetles, earwigs and isopods (seen eating seeds during depot checks; ants and slugs also readily consume sunflower seeds during feeding trials) (*12, 37*). Sunflower seeds therefore measure total predation by the full suite of seed-predator groups. To measure invertebrate-only predation, we excluded vertebrates from some sunflower depots using cages of ½” (∼1.3 cm) mesh (Fig. S1).

In contrast, carbohydrate-based oat seeds are eaten almost exclusively by small mammals (detected via feces left in depots) and birds (camera traps). Across 5376 oat depots deployed here and in (*12*), only 1 of 26,880 oat seeds was eaten by ants (1 ant found eating a seed at 24 h check; other 4 oats intact). Mollusc predation on oats was seen twice, and isopod predation seen once. In all cases, invertebrates had made only small holes in oats after 24 hours, and no oats had been entirely eaten or removed, so it seems unlikely that we are missing frequent oat predation by invertebrates. Predation on oats thus reflects almost-entirely vertebrate predation.

After 24 h we searched for seed remains in and around each depot. We counted seeds whose endosperm was partially eaten, fully eaten (but with sunflower seed skins or oat hulls remaining), or removed. We noted signs of predators, e.g. rodent feces, slug slime trails, invertebrates still eating seeds. Seeds removed from the ground can occasionally be dispersed rather than eaten, known as secondary dispersal. Nevertheless, we counted missing seeds as eaten for three reasons: 1) more than 90% of seeds removed from the ground are eaten (*55*); 2) our seeds have no adaptations to facilitate secondary dispersal (e.g. no thick shells or elaiosomes) making secondary dispersal especially unlikely; and 3) we know from observations and camera trap footage that many of our major predator groups remove both sunflower and oat seeds and consume them immediately, further suggesting that most removed seeds are eaten rather than dispersed. All birds captured on camera traps (jays, robins, sparrows) consumed seeds immediately. Ants were often seen removing seeds from depots but were almost always breaking seeds up and eating them at the same time.

#### Identifying seed predators

In most cases of seed predation, there were no definitive signs of the seed predator found (4356 of 5825 depots with predation). However, for about a quarter of depots with predation (1469 depots), we were able to identify seed predators as follows.

<u>Ant</u> predation was noted during depot checks if ants were seen eating seeds (either in depots or while carrying seeds away from depots), if ant hills had been built inside the depot (usually in cages), or if all seeds had been removed but ants were still swarming the depot.

<u>Mollusc</u> predation was noted if snails or slugs were seen eating seeds or if seeds had been eaten and there were slime trails through the depot.

<u>Mammals</u> were never seen directly, but sometimes left tell-tale signs in depots. Small mammals (e.g. mice, voles) sometimes left feces in depots. When feces were found in oat depots, we often also found the thin hulls of oat seeds left behind; we therefore considered depots in which oats had been eaten but hulls left behind to be predated by small mammals. Finally if cages were dug up or under, or if seeds had been eaten and the wooden popsicle stick marking the depot was chewed with visible incisor marks, we counted that as signs of mammal predation.

<u>Other</u> predator types (e.g. birds caught on camera traps, beetles or other invertebrates seen eating seeds) were recorded but were rare.

These observations are useful for comparison among sites, as we do below, but should be interpreted with nuance. First, ant predation usually needed to be directly observed, so in climates or sites where ants were particularly active and removed all seeds early on, ant predation would be underestimated. Second, predator groups can affect each other. For example, it is only possible to see ant predation if vertebrates have not already eaten seeds and vice versa. When comparing predator observations among sites, we took these constraints into account as follows.

- Models of ant predation: considered only caged depots, which measure invertebrate predation rates independent of vertebrate predation. Cages also made it more difficult for ants to completely remove seeds, meaning we were more likely to see ant predation still in action after 24 h.
- Models of small mammal predation considered only oats, as a) oats were almost never eaten by invertebrates and so provide a measure of mammal predation independent of invertebrate predation and b) mammals often peeled hulls from oat seeds and left them in the depot, so we were better able to detect mammal predation on oats than on sunflower seeds which were eaten whole.
- Models of mollusc predation considered all depot types, as molluscs often left slime trails so were easily observed in both caged and uncaged depots. We included oat depots, as molluscs were twice seen eating oats (of > 5000 oat depots). More often molluscs left slime trails in oat depots without eating seeds (in which case depots were not recorded as predated). Results do not change if we consider only caged sunflower depots.

**Fig. S2.**
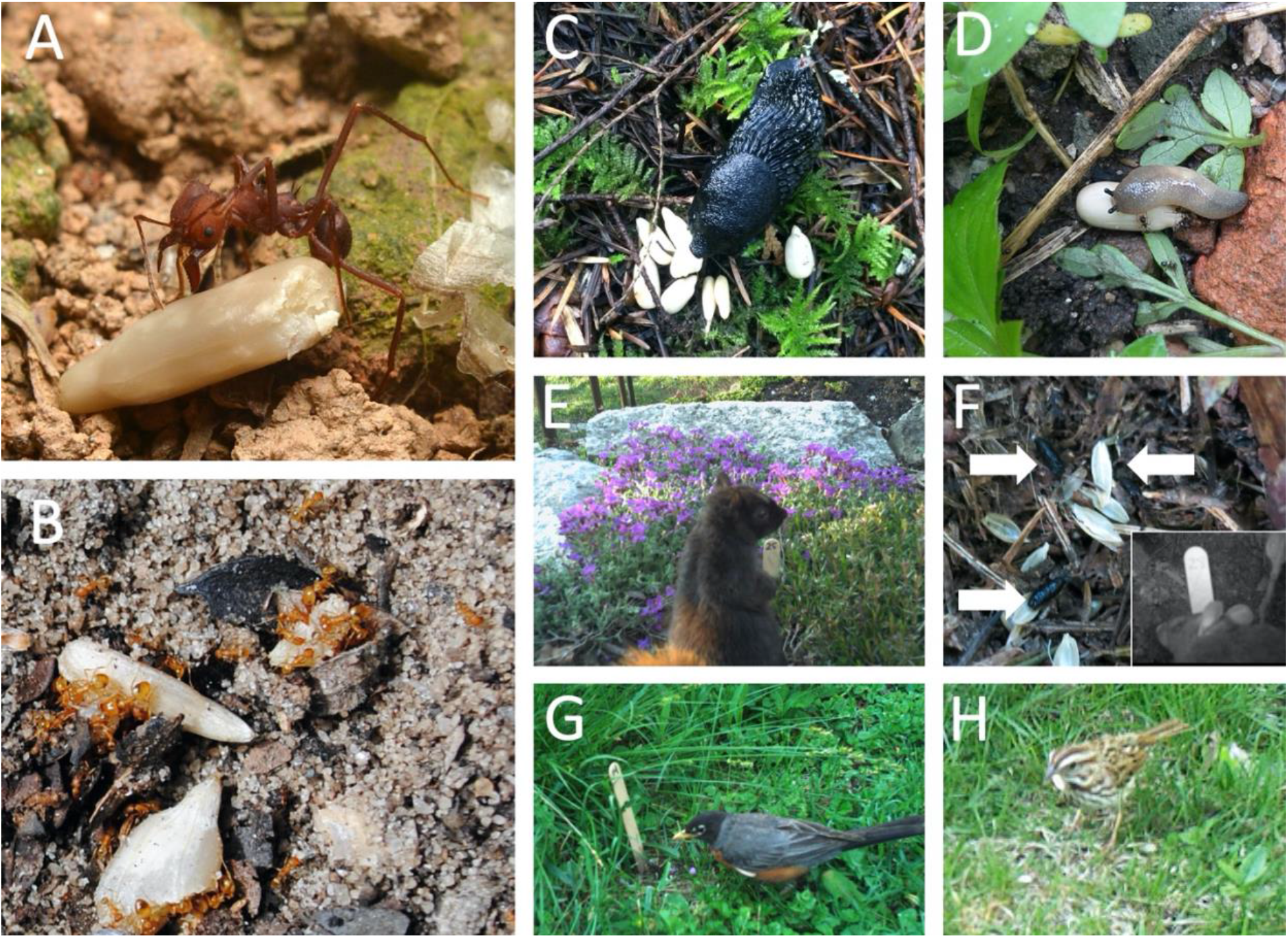
Seed predators. Ants (A-B) and molluscs (C-D) were the most frequently detected invertebrate seed predators. Both molluscs and ants sometimes left the papery coat of sunflower seeds behind (A, on right). E) Squirrels and F) small rodents (rats, mice) ate both sunflower and oat seeds (caught on camera traps). F) Small rodent predation was also evident from droppings left in depots (right pointing arrows) and, for oats, thin hulls carefully stripped from seeds (left pointing arrow). G-H) A variety of birds were caught on camera traps eating both sunflower seeds (robins, jays, sparrows, finches) and oats (sparrows). Photo credits: A=L Paolucci (Vicosa, Brazil); B=F Oliveira (Buique, Brazil); C=T Veen (Squamish, Canada); D=C García Jiminez (Guadalajara, Mexico); EG=A Hargreaves (eastern grey squirrel and American robin eating sunflower seeds, Kingston Canada); F=A Hargreaves (Vancouver, Canada), F inset =J Muñoz (rodent eating sunflower seeds, Colombia); H=A Hargreaves (song sparrow eating oat, Kingston Canada).

#### Covariates

Urbanized sites varied greatly in how natural they were and how urbanized their neighbourhoods were, and local natural areas also varied in size and connectivity. We therefore investigated covariates that might explain seed predation rates better than our simple categorization of urbanized or natural. First, during experiments we recorded the ground *vegetation type* surrounding each depot (i.e. a microsite): lawn (primarily grass that is regularly mowed); garden (planted and tended areas with plants surrounded by bare soil, mulch, or non-grass ground cover); or untended areas with natural vegetation litter (whether or not the litter was from native plants). At each site we set depots out to sample each habitat type roughly in proportion to its area (e.g. if a backyard had 50% lawn and 50% garden we set out roughly half our depots in each vegetation type). All depots in natural sites were in areas with untended, natural vegetation litter.

Second, we scored the greenness at each site, using satellite images (GoogleEarth) processed by hand using our local knowledge of each site, rather than relying on interpolated rasterized databases. We calculated the *% green area within 500 m* (i.e. a 500 m radius from the centre of each experimental site; ImageJ). We first cropped out water bodies (lakes, large rivers). We then quantified the green area, defined as having vegetated ground cover, i.e. including gardens, lawns, fields, and natural vegetation and excluding ‘grey’ areas defined as those covered with built structures, paving, stonework or gravel. Areas with green canopy but grey ground covering (e.g. tree canopies extending over roads) were counted as grey. We also calculated the *connected green area* (m^2^) for each site, as the total green area that could be accessed without crossing grey areas (ignoring fences). This was done using GoogleEarth’s polygon feature. GoogleEarth images were downloaded within 6 months of the final experimental run at each site.

#### Statistical analyses

Analyses were performed in R (v 4.0.2) (*56*). Analyses of seed predation (proportion seeds eaten) used generalized linear mixed models (GLMMs) with a binomial error distribution and logit-link. We had 1 data point per depot, and models included random intercepts for run and site, and an individual-level random intercept (depot) to resolve overdispersion (*57*). We tested the significance of model fixed effects, beginning with higher order interactions, using likelihood ratio tests, comparing the resulting ratios to a Chi-squared distribution. When interactions were significant we assessed which trends or means differed using estimated marginal means (*58*). For visualization, we extracted trendlines and partial residuals from models (*59*), and estimated 95% confidence intervals following (*57*).

Our main analysis asked whether seed predation varied with latitude for total, invertebrate, or vertebrate predation. Our full model had fixed effects Latitude × Urbanization × Predation type + Elevation. Latitude is absolute latitude in decimal degrees, Urbanization is categorical (urbanized or natural), and Predation is categorical, as measured by our three depot types (uncaged sunflower, caged sunflower, or oat, measuring total, invertebrate, or vertebrate predation respectively). We did not choose sites to vary systematically in elevation, so we simply controlled for its effect as seed predation varies strongly with elevation (*12, 60*). Our full data set included 337 experimental runs at 81 sites. We explored the robustness of our results (see Table S2, Fig. S2 for full results). First, we re-ran the overall model including only runs where an urbanized site was tested within 2 weeks of a paired natural site (303 runs at 74 sites). Second, we ran one model per predation type with fixed effects Latitude × Urbanization + Elevation.

We next explored the effects of surrounding ground vegetation and greenness. We first tested whether our three measures of local vegetation (proportion of depots in microsites with unkept vegetation with natural litter, % green area within a 500 m radius, total connected green area), varied between urbanized and natural sites or with latitude (full statistical details and results in Table S1). Second, we tested whether either measure of greenness explained seed predation better than simply classifying a site as urbanized or natural, by replacing the Urbanization term in our main analyses with either greenness measure (scaled). The combined effect of greenness and latitude differed among predation types (%Greenness × Latitude × Predation type: χ^2^_df=2_ = 7.41, *P* = 0.025; Connected green area × Latitude × Predation type: χ^2^_df=2_ = 63.39, *P* < 0.0001), so we then ran one model per predation type with fixed effects Greenness × Latitude + Elevation.

Data and code will be publicly archived upon publication.

### Supplemental Results

**Table S1.** Ground vegetation and local greenness in natural vs. urbanized sites. We tested whether each response variable differed between urbanized and natural sites, and whether this difference varied with latitude (model fixed effects = Urbanization (categorical; urbanized or natural), Latitude (absolute), and their interaction). For the proportion of seed depots placed in microsites with natural vegetation and litter, we ran a GLMM with a random intercept for site. For greenness analyses, model fixed effects were Urbanization, Latitude, and their interaction. To improve model fit, connected green area was log transformed. Significance of fixed effects was determined using likelihood ratio tests; for GLMMs and linear models the resulting ratios are compared to a Chi-squared or *F*-distribution, respectively. Significance of 2-way interaction compares the full model to a model without the interaction. As the interaction was never significant, it was dropped, and significance of individual terms is comparing a model without the term of interest to the reduced (no interaction) model. A significant effect of Urbanization confirms that sites in urbanized areas differ categorically from those in natural areas. The Latitude × Urbanization interaction tests whether urbanized and natural sites become more similar at one end of the latitudinal gradient, and the Latitude term tests whether the degree of modification varies with latitude, either of which could bias our detection of latitudinal patterns. Results of the Urbanization comparison are plotted in Fig 1C.

| Response | Response structure | Model type | Latitude x Urbanization | Urbanization | Latitude |
| --- | --- | --- | --- | --- | --- |
| depot in untended, natural vegetation or not | Binary, 1 data point per depot | Binomial GLMM | $\chi^2_{df=1} < 0.01$<br>$P > 0.99$ | $\chi^2_{df=1} = 114.1$<br>$P < \mathbf{0.0001}$ | $\chi^2_{df=1} = 0.16$<br>$P = 0.69$ |
| % green area within a 500 m radius | %<br>1 data point per site | Linear | $F_{df=1} = 0.05$<br>$P = 0.82$ | $F_{df=1} = 26.3$<br>$P < \mathbf{0.0001}$ | $F_{df=1} = 0.3$<br>$P = 0.57$ |
| Connected green area (m<sup>2</sup>) | Log(m<sup>2</sup>) outliers constrained<sup>1</sup><br>1 data point per site | Linear | $F_{df=1} = 0.46$<br>$P = 0.50$ | $F_{df=1} = 104.7$<br>$P < \mathbf{0.0001}$ | $F_{df=1} = 0.1$<br>$P = 0.78$ |

**Fig. S3.**
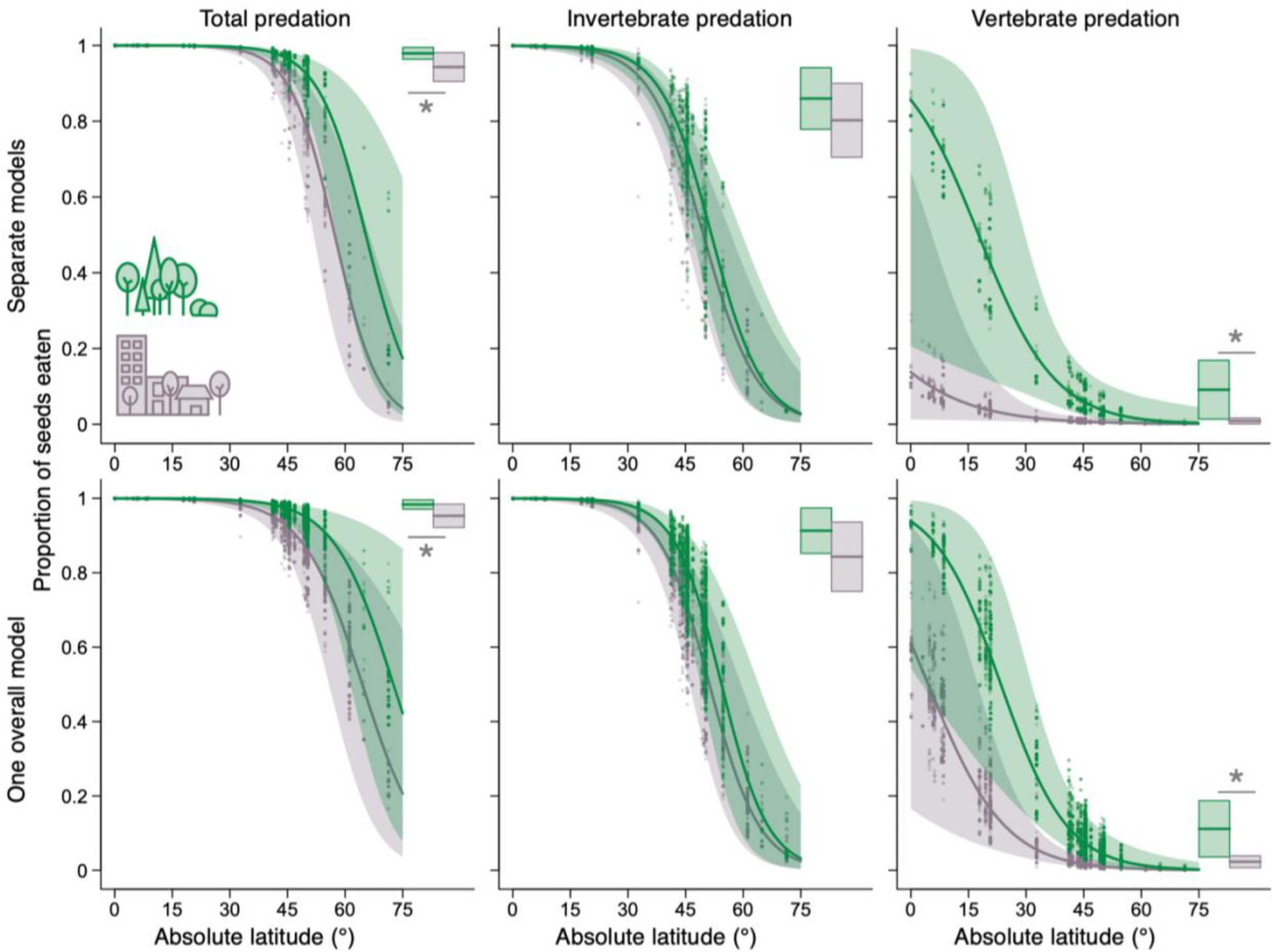
Separate models per predation types vs one overall model for all predation types. Top row is the same as Fig. 2, and shows results from 1 model per predation type, each with fixed effects Latitude × Urbanization + Elevation. Bottom row shows results from a single overall model including all predation types, with fixed effects Latitude × Urbanization × Predation type + Elevation. Overall conclusions are the same (full statistical results in Table S1). Thick lines, shaded polygons and points show means, 95% CI, and partial residuals, respectively.

**Table S2.** Statistical results were consistent across data structures, whether models included: data from all runs and all predation types; data from all predation types but only paired runs (i.e. excluding runs when an urbanized site was tested without a concurrent test in a paired natural area); or only one predation type at a time. A ‘run’ is one 24 h assay of predation at one site. The only qualitative difference is in how much urbanization reduced total predation (C).

**A) Sample sizes.**
| Data | Site type | Sites | Runs |
| --- | --- | --- | --- |
| All runs | Urbanized | 45 | 181 |
|  | Natural | 36 | 156 |
|  | Total | 81 | 337 |
| Paired runs only | Urbanized | 38 | 148 |
|  | Natural | 36 | 155 |
|  | Total | 74 | 303 |

|  |  | Results of Likelihood Ratio tests |  |  |  |  |
| --- | --- | --- | --- | --- | --- | --- |
| Data | Predation type | Urban × Lat × Pred type | Urban × Lat | Urban × Pred type | Lat × Pred type | Elevation |
| All runs | All | $\chi^2_{df=2} = 1.17$<br>$P = 0.55$ | $\chi^2_{df=1} = 0.11$<br>$P = 0.74$ | $\chi^2_{df=2} = 17.85$<br>**$P = 0.0001$** | $\chi^2_{df=2} = 54.11$<br>**$P < 0.0001$** | $\chi^2_{df=1} = 26.59$<br>**$P < 0.0001$** |
| Paired runs only | All | $\chi^2_{df=2} < 0.01$<br>$P > 0.99$ | $\chi^2_{df=1} = 0.19$<br>$P = 0.67$ | $\chi^2_{df=2} = 41.52$<br>**$P < 0.0001$** | $\chi^2_{df=2} = 71.31$<br>**$P < 0.0001$** | $\chi^2_{df=1} = 19.88$<br>**$P < 0.0001$** |
| All runs | Invertebrate | – | $\chi^2_{df=1} = 0.07$<br>$P = 0.79$ | – | – | $\chi^2_{df=1} = 25.67$<br>**$P < 0.0001$** |
| All runs | Vertebrate | – | $\chi^2_{df=1} = 0.19$<br>$P = 0.66$ | – | – | $\chi^2_{df=1} = 9.41$<br>**$P = 0.0022$** |
| All runs | Total | – | $\chi^2_{df=1} = 0.19$<br>$P = 0.66$ | – | – | $\chi^2_{df=1} = 43.51$<br>**$P < 0.0001$** |

| Data (model) structure | Effect of Urbanization × Predation type | | Effect of Latitude × Predation type | |
| --- | --- | --- | --- | --- |
|  | Least-squares means contrasts urbanized vs. natural sites | Where is predation highest? | Latitudinal trend significant? (95% CI don't overlap 0) | Differences in Latitudinal trend among Predation types |
| All runs (1 overall model) |  |  |  |  |
| Total predation | $z = 2.1$<br>**$P = 0.0395$** | natural > urban | **Yes** | b |
| Invertebrate | $z = 1.3$<br>$P = 0.21$ | — | **Yes** | a (steepest) |
| Vertebrate | $z = 3.2$<br>**$P = 0.0014$** | natural > urban | **Yes** | ab |
| Paired runs (1 overall model) |  |  |  |  |
| Total predation | $z = 1.1$<br>$P = 0.27$ | — | **Yes** | b |
| Invertebrate | $z = 0.2$<br>$P = 0.82$ | — | **Yes** | a (steepest) |
| Vertebrate | $z = 3.1$<br>**$P = 0.0021$** | natural > urban | **Yes** | ab |
| All runs (1 model per predation type) |  |  |  |  |
|  | LRtest for Urbanization |  | LRtest for Latitude |  |
| Invertebrate | $\chi^2 = 0.73$<br>$P = 0.39$ | — | $\chi^2 = 34.28$<br>**$P < 0.0001$** | — |
| Vertebrate | $\chi^2 = 4.07$<br>**$P = 0.044$** | natural > urban | $\chi^2 = 7.77$<br>**$P = 0.0053$** | — |
| Total predation | $\chi^2 = 4.07$<br>**$P = 0.044$** | natural > urban | $\chi^2 = 40.09$<br>**$P < 0.0001$** | — |

**Table S3.** Seed predator observations varied with urbanization and latitude. For our three most commonly detected seed predators, we tested whether they were detected more often in natural or urbanized sites and whether this varied with latitude (one binomial GLMM per predator). Full model fixed effects were Urbanization × Latitude + Elevation, with a random intercept for site. Likelihood ratio tests first compare models with and without the 2-way interaction; as it was not significant it was dropped and subsequent tests compare models without the fixed effect of interest to the no-interaction model. Ant models consider only predation from caged depots, because caged depots measure invertebrate predation independent of vertebrate predation rates. Further, cages slow seed removal by ants, making it more likely ant predation can still be detected after 24 h (vs. if all seeds are entirely removed well before depots are checked). Models of small mammal predation consider only predation on oat depots, as oats are almost entirely ignored by invertebrates and so measure vertebrate predation independent of invertebrate predation. Further, as rodents often strip hulls from oats and leave them in the depot, small mammal predation is easier to detect for oat vs. sunflower seeds. Mollusc presence in depots is easily detected even if all seeds have been eaten, as snails and slugs leave visible slime trails; mollusc models therefore consider all seed types (results are consistent if we consider only caged depots).

| Seed predator | Total depots at which predation detected | Depot types included in model | Results of Likelihood ratio tests |  |  |  |
| --- | --- | --- | --- | --- | --- | --- |
| | | | Lat × Urbanization (df = 1) | Urbanization (df = 1) | Latitude (df = 1) | Elevation (df=1) |
| Ants | 685 | Caged sunflower | $\chi^2 = 0.10$<br>$P = 0.75$ | $\chi^2 = 13.12$<br>**$P = 0.0003$** | $\chi^2 = 4.74$<br>**$P = 0.029$** | $\chi^2 = 5.38$<br>**$P = 0.020$** |
| Mammal | 538 | Oats | $\chi^2 = 0.33$<br>$P = 0.57$ | $\chi^2 = 15.34$<br>**$P < 0.0001$** | $\chi^2 = 0.20$<br>$P = 0.66$ | $\chi^2 = 0.48$<br>$P = 0.49$ |
| Mollusc | 263 | All | $\chi^2 = 0.19$<br>$P = 0.66$ | $\chi^2 = 2.80$<br>$P = 0.094$ | $\chi^2 = 2.58$<br>$P = 0.11$ | $\chi^2 = 0.48$<br>$P = 0.49$ |
| Other | 54 | — | — | — | — | — |

**Table S4.** Relative fit and explanatory power of different measures of greenness. All models are binomial GLMMs, 1 per predation type. Full model fixed effects are absolute Latitude × Greenness measure. Models include random intercepts for run (i.e. date within site), site and observation, except the invertebrate model for % greenspace, in which the random intercept for run is replaced with a simpler random intercept for date (not nested within site) to improve convergence. Significance of terms is determined using likelihood ratio tests (comparing models with fixed effects ‘Greenness × Lat’ to ‘Greenness + Lat’, or ‘Greenness + Lat’ to ‘Lat’). Δ values compare greenness model to the equivalent (same random effects structure) model with categorical Urbanization as the fixed effect, with Δs calculated such that negative values mean the greenness models perform worse (i.e. have higher AIC or lower R^2^_GLMM_). i.e. ΔAIC = AIC_cat.urban_ – AIC_greenness_, and ΔR^2^_GLMM_ = R^2^_greenness_ – R^2^_cat.urban_. R^2^_GLMM_ is the marginal (pseudo) R^2^ for GLMMs, which shows the relative explanatory power of model fixed effects (*61*). Binomial variance is calculated as the observation-level variance (delta method in the MuMIn package (*62*)).

| Predation<br>type | Greenness = % greenspace within 500 m |  |  |  | Greenness = connected green area (m<sup>2</sup>) |  |  |  |
| --- | --- | --- | --- | --- | --- | --- | --- | --- |
| | Greenness<br>× Lat | Greenness | $\Delta\text{AIC}$ | $\Delta R^2_{\text{GLMM}}$ | Greenness<br>× Lat | Greenness | $\Delta\text{AIC}$ | $\Delta R^2_{\text{GLMM}}$ |
| Total | $\chi^2 = 0.57$<br>$P = 0.45$ | $\chi^2 = 1.24$<br>$P = 0.27$ | -2.8 | -0.0080 | $\chi^2 = 0.91$<br>$P = 0.34$ | $\chi^2 = 0.90$<br>$P = 0.34$ | -3.2 | -0.0037 |
| Invert. | $\chi^2 = 0.17$<br>$P = 0.68$ | $\chi^2 = 3.49$<br>$P = 0.062$ | 3.0 | 0.0052 | $\chi^2 = 4.09^*$<br>**$P = 0.043$** | – | 1.9 | 0.0361 |
| Vert. | $\chi^2 = 1.11$<br>$P = 0.29$ | $\chi^2 = 0.03$<br>$P = 0.86$ | -13.5 | -0.0338 | $\chi^2 = 0.76$<br>$P = 0.38$ | $\chi^2 = 0.67$<br>$P = 0.41$ | -12.9 | -0.0315 |

**Table S5.**
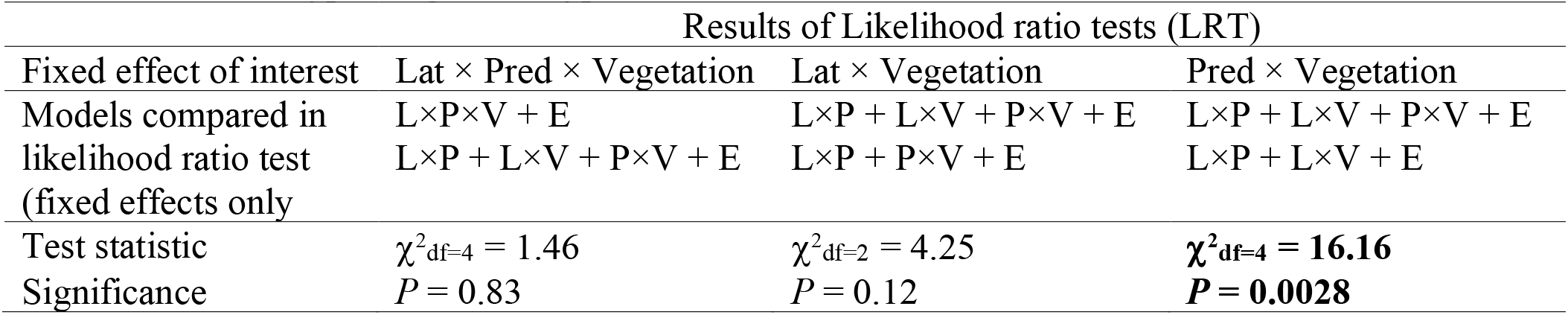
Effect of ground vegetation type on predation in urbanized sites. Vegetation type is where each depot was placed = lawn (grass and other herbaceous cover, regularly mowed), garden (planted beds, usually with ground cover of bare soil or mulch), or natural (natural, untended vegetation and litter, whether or not plants are native). Model fixed effects are absolute Latitude, Predation type, Vegetation type, and Elevation.

**Fig. S4.**
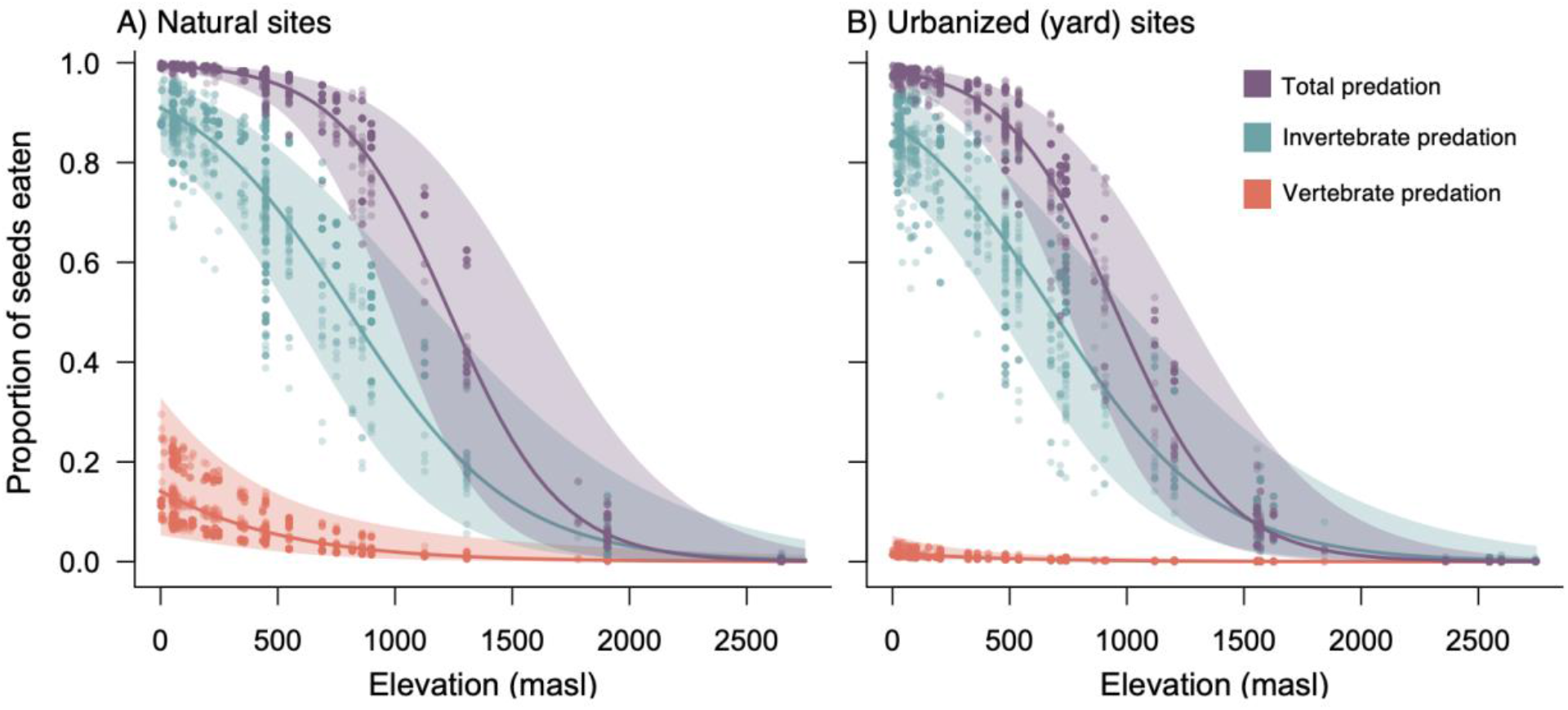
Seed predation declined toward higher elevations. While our study was not designed to test for elevational patterns in seed predation (unlike (*12*), which systematically sampled across elevations within each latitude), we included elevation as a covariate in all analyses of seed predation to account for its effects, and it was always significant (Table S2). Elevational trends (± 95% CI) and partial residuals are extracted from one GLMM per predation type (fixed effects: Latitude (absolute) × Urbanization (categorical) + Elevation), and are plotted separately for natural and urbanized sites for illustration. Statistical results in Table S1B).

## Notes

### Competing Interest Statement

The authors have declared no competing interest.

